# Polymer Implantable Electrode Foundry: A shared resource for manufacturing polymer-based microelectrodes for neural interfaces

**DOI:** 10.1101/2023.11.05.565048

**Authors:** Kee Scholten, Huijing Xu, Zhouxiao Lu, Wenxuan Jiang, Jessica Ortigoza-Diaz, Artin Petrossians, Steven Orler, Rachael Gallonio, Xin Liu, Dong Song, Ellis Meng

## Abstract

Large scale monitoring of neural activity at the single unit level can be achieved via electrophysiological recording using implanted microelectrodes. While neuroscience researchers have widely employed chronically implanted electrode-based interfaces for this purpose, a commonly encountered limitation is loss of highly resolved signals arising from immunological response over time. Next generation electrode-based interfaces improve longitudinal signal quality using the strategy of stabilizing the device-tissue interface with microelectrode arrays constructed from soft and flexible polymer materials. The limited availability of such polymer microelectrode arrays has restricted access to a small number of researchers able to build their own custom devices or who have developed specific collaborations with engineering researchers who can produce them. Here, a new technology resource model is introduced that seeks to widely increase access to polymer microelectrode arrays by the neuroscience research community. The Polymer Implantable Electrode (PIE) Foundry provides custom and standardized polymer microelectrode arrays as well as training and guidance on best-practices for implantation and chronic experiments.

## 1. Introduction

Chronically viable neuroelectronic interfaces are required for neuromodulation therapy and fundamental neuroscience research, but many available tools suffer from biological failure modes due to the mechanical stress and damage inflicted on neural tissue following implantation [1–6]. Most notably penetrative neural implants targeting the brain, historically made from rigid materials such as silicon and metal, struggle to reliably track specific neuronal units across chronic animal studies [1,7]. With regards to applications targeting peripheral nerves, such stiff materials lack the mechanical flexibility and conformality needed to match the complicated contours of peripheral nerve anatomy. These limitations have driven the development of polymer microelectrode arrays (pMEAs), an emerging technology that can replace silicon and metal-based electrodes with softer and more flexible alternatives as a means to mitigate the immune response and expand the range of accessible target anatomies.

An extensive number and range of pMEAs have been described in the literature, customized to the anatomical requirements and size scales of different species from invertebrates, including hookworm [8] and earthworm [9], to vertebrates, including zebrafish [10], salamander [11], mouse [12], rat [13–16], dog [12,17], and monkey [18,19]. In recent years several efforts have led to the creation of highly sophisticated pMEAs, featuring hundreds to thousands of electrodes integrated with application specific circuitry for on-site multiplexing or signal modulation [6,20,21]. To date, however, pMEAs remain a niche technology among neuroscientists, and among those using pMEAs, there is little standardization or consolidation among the various materials, designs, and approaches, making rigorous comparison between studies difficult.

This is attributed to the difficulty in accessing pMEAs which are not available off-the-shelf as a commercial technology, except for a limited selection of cuff electrodes and surface arrays (e.g., arrays for electroencephalogram (EEG) and electrocorticography (ECoG)). Customized pMEAs are instead produced via collaborations supported by sponsored research between neuroscientists and select academic laboratories with both microelectronics expertise and access to micro/nanofabrication facilities. Since such collaborations usually target a specific region in a specific species, this one-by-one approach yields only a few custom-designed devices suitable for initial demonstration purposes but typically does not result in device production at scale to support larger and repetitive research studies. This is exacerbated by the reliance on research trainees to design, fabricate, and package devices as an intermediate step in their career development and the absence of affordable and convenient contract research services to develop custom microelectrode arrays. Although some startup companies have resulted from such efforts, this avenue of access does not guarantee a sustained supply of custom pMEA designs.

In many ways, this technological trajectory mirrors the history of conventional microelectrode arrays (MEAs). Academic researchers developed the first MEAs to advance fundamental neuroscience research; early devices were manually produced from hand-wound microwire bundles [22–24], which later evolved into silicon-based arrays produced using photolithography and semiconductor micromachining as part of student directed research [25–29]. After many iterations of development led to increased channel count and complexity, commercial efforts arose to help disseminate standardized forms of these silicon MEAs to researchers across the globe. The successful transition from bespoke laboratory device to mass-produced commercial offering was driven by key manufacturing intermediaries, and in particular the Metal Oxide Semiconductor Implementation Service (MOSIS) [30].

MOSIS is a low-cost integrated circuit prototyping service which helps provide manufacturing of semiconductor devices to university researchers and students. Launched with US government funding in the 1980s, MOSIS transitioned to a service fee model, and has since supported 50 US government laboratories and agencies, 800 colleges and universities worldwide, and over 100 companies [31]. MOSIS operates by consolidating different projects from different users onto a single wafer, creating a multi-project wafer (MPW); devices are then manufactured simultaneously using a standardized set of processes, distributing costs across all projects such that the small-volume manufacturing typical for research becomes affordable. MOSIS played a vital role in the development of silicon MEAs, providing manufacturing of electrodes [32–36] and ancillary circuit components [37–41] at a quality and cost that could not be reliably produced by academic labs or start-ups, at a time before serious commercial efforts were available. The shared-resource model demonstrated by MOSIS was a necessary steppingstone in the maturation of silicon MEAs as a technology.

MOSIS leverages the large semiconductor manufacturing industry and established silicon foundries, taking advantage of half a century of investment and research into silicon micromachining techniques. But polymer micromachining remains a nascent practice; commercial foundries permit few if any polymer substrates in their process flows, and those are often limited to polyimide for the production of flexible PCBs or cables. For researchers who require pMEA technology for neuroelectronic interfaces and lack the facilities or training to build their own, there are few options. Commercial pMEAs are available in a limited number of designs, necessarily driven by the need to scale production and achieve profits. However, researchers require not only access to a sufficient quantity of high-quality devices, but pMEAs of custom designs to meet the specific needs of their target animal model, anatomy, and experiments. For example, multi-shank pMEAs with specifically determined inter-shank spacings may be needed to span multiple cortical regions, conformally arranged electrodes are needed to efficiently record neural activities from multiple hippocampal sub-regions simultaneously, and surface electrodes in a custom-fit, extrafasicular cuff design, are needed to target peripheral nerves with precise diameters. As such, the neuroscience community needs stable and easy access to design services and the manufacturing capabilities that drove silicon MEA proliferation.

To address the needs of neuroscience and neural engineering researchers we launched a shared technology resource center dubbed the Polymer Implantable Electrode (PIE) Foundry. The PIE Foundry is a professionally staffed service center that offers manufacturing, packaging, testing, support, and dissemination of customized pMEAs for research, using a standardized set of fabrication protocols to create reliable and comparable devices, currently offered at no cost to the academic and nonprofit research community (users). In addition, the PIE Foundry offers virtual and on-site training services on pMEA design, packaging, testing, and implementation in animal studies. Polymer microfabrication is performed in cleanroom facilities and laboratories at the University of Southern California, using a set of well-established polymer microlithographic processes developed for the creation of implantable biomedical microdevices [42–45]. Like MOSIS, the PIE Foundry operates on an MPW model; multiple user designs are manufactured simultaneously on a shared silicon wafer using processes capable of producing pMEAs of arbitrary design, size, channel count, and form-factor. The PIE Foundry goes beyond the services offered by MOSIS as it can directly fabricate user-submitted designs as well as allowing users to request available designs from a small library of existing pMEAs. In addition, for users requiring designs that deviate from the standard capabilities, materials, or other design rules, a small number of custom projects, based on a single project wafer (SPW) model, are offered each year; here, the entire wafer is dedicated to a single user. Such custom projects require the approval of an independent steering committee which evaluates the proposal for utilization of PIE Foundry resources, thereby providing independent review of access to the resource.

The PIE Foundry is currently supported as a research resource grant (U24 NS113647) through the National Institutes of Health (NIH) under the BRAIN Initiative [46], which allows research user access to services at no cost and at cost to commercial users. Principal operation consists of manufacturing and testing pMEAs for academic users at small-batch scale, typically in batches of 10-100 individual dies. These pMEAs span a large range of designs and applications, from peripheral nerve stimulators to high-density surface or penetrating recording arrays. In all cases, designs are intended for *in vivo* experiments in animal models, and examples of models investigated by users span songbird, to rat, to sheep.

Currently, the PIE Foundry uses Parylene C as both the polymer base and electrical insulation for all pMEAs. There are several options for polymer materials used in neural interfaces, the most popular and mature being polyimide [6,20,47–49] and Parylene C [13,14,55,56,15–17,50–54]. Both have similar Young’s moduli, densities, and electrical properties, and an extensive recent history in neural interface development [57]; however, we selected Parylene C due to its high optical transparency, useful thermoplastic properties, long history in FDA approved implanted devices, unique ability to be coated using room temperature vapor deposition, and our extensive prior experience in Parylene C micromachining [42,45]. The most common structure of a PIE Foundry pMEA comprises a symmetric polymer-metal-polymer sandwich, with a Parylene C base supporting a thin-film metal layer containing electrodes, traces, and contact pads, and a final Parylene insulation layer. Oxygen plasma is used to selectively remove polymer, exposing contacts and electrodes, and cutting out the Parylene C into arbitrary two-dimensional shapes, including penetrating probes, surface recording arrays, and peripheral nerve interfaces including spinal arrays (Fig. 1). While simple, this approach is incredibly flexible; pMEAs can be built having nearly arbitrary channel count and any size or shape, and using the MPW model, simultaneous microfabrication of varied and distinct devices having nearly identical properties is achieved by using shared materials and processes.

**Figure 1.**
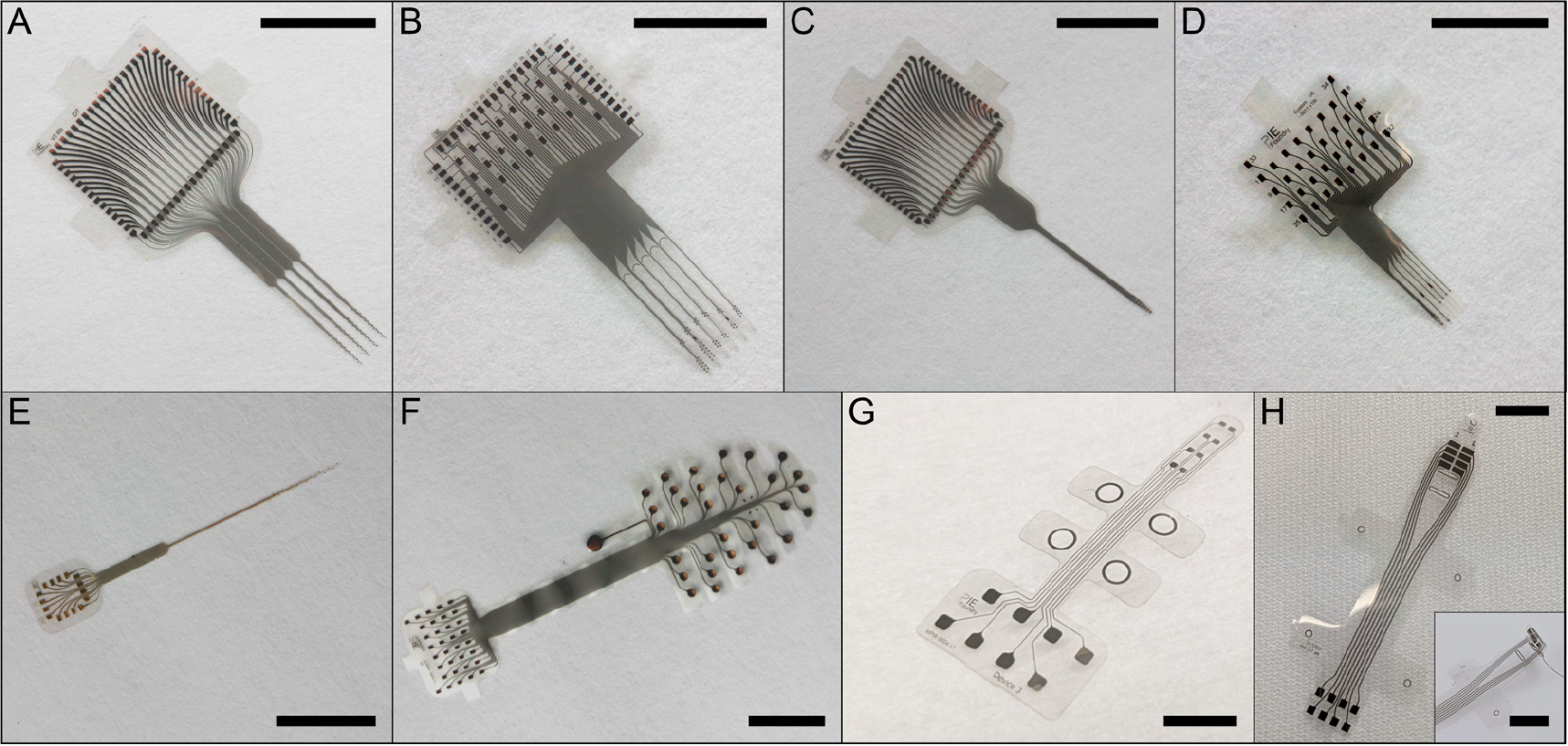
Photographs of polymer microelectrode arrays manufactured by the PIE Foundry for external users in a variety of sizes, shapes, and form-factors. Scalebars are 5 mm. **A**: The PIE Foundry ‘standard’ array, a 64-channel, 4-shank penetrating array for (sub)cortical recording in small mammals. **B**: 64-channel, 6-shank penetrating array for targeted hippocampal recording in rats. **C**: 32-channel, single shank penetrating tetrode array for spike detection and sorting in small mammals. **D:** 32-channel, 4-shank penetrating array for targeted hippocampal recording in mice. **E**: 16-channel, gold coated penetrating array for deep brain recording in cats. **F**: 32-channel surface array for electroencephalography in small mammals. **G**: 8-channel paddle electrode array for stimulating the spinal cord in small mammals; the design is attributed to the Grill Lab at Duke University. **H**: 8-channel peripheral nerve array for recording and simulation from peripheral nerves; design is attributed to the Bruns Lab at the University of Michigan. Inset shows the array in a curled cuff configuration.

Here we detail the methods and outcomes of the PIE Foundry operations from 2019-2023, including key results of the shared-resource model, in-depth polymer microfabrication process steps, and representative data from select *in vivo* experiments.

## 2. Materials & methods

### 2.1. Shared-resource model

The PIE Foundry uses several research core facilities (cores for nanofabrication, electron microscopy, and conventional machining) and the labs of Dong Song and Ellis Meng at the University of Southern California. Full-time professional staff support the design, fabrication, and testing of pMEAs requested by the neuroscience and neural engineering research community. In addition to providing electrode arrays, users have access to training, including multielectrode array design, fabrication protocols, and the implementation of flexible electrodes in animal studies.

There are three distinct routes that have evolved since the founding of the PIE Foundry for users to obtain pMEAs. Users proficient in computer aided design tools can generate and submit their own designs in a standard format (.gbr, .dwg, .dxf, .gds) which are then pooled with those of other users for simultaneous fabrication as part of the MPW model. This requires that designs adhere to a set of guidelines to facilitate packing of multiple projects into distinct regions on the same carrier wafer. These designs are all processed together using the same microfabrication recipe and so the materials are identical and only the designs differ.

If a user project requires non-standard materials or processing steps, users can submit a request for a custom, single-project wafer fabrication run. Examples of non-standard requirements include user-specified materials for electrodes, multi-layer fabrication including the use of connective vias, or features beyond the typical pattern resolution of an MPW run. This avenue is also relevant for users who have defined a set of requirements for an electrode array but lack the expertise in device layout. Since significant staff time or a change in process are required, there is added effort and cost beyond the standard process offered. An external scientific steering group is retained to review submission of requests provided in the form of a short proposal. This review mechanism ensures that efforts undertaken are scientifically significant and within the resource’s mission while also providing a mechanism to manage any potential conflict of interest. Approved projects receive PIE Foundry staff support for design, fabrication, packaging, and basic characterization of produced pMEAs. Table 1 compares the offered capabilities for MPW processing with SPW processing for custom projects.

**Table 1:** Parameters of manufacturing capabilities for single- and multi-project wafer processing.

| Capabilities |  | Multi-Project Wafer | Single-Project Wafer |
| --- | --- | --- | --- |
| **Material** | Insulation (polymer) | Parylene C | Parylene C |
|  | Conductor (metal) | Pt or Ti/Pt/Au/Pt | Ti, Au, Pt, Cr, Al |
| **Layer thickness** | Insulation (polymer) | 10 $\mu\text{m}$ | 2 -20 $\mu\text{m}$ |
|  | Conductor (metal) | 200 nm | 50 – 500 nm |
| **Minimum pattern resolution** | Insulation (polymer) | 10 $\mu\text{m}$ | 5 $\mu\text{m}$ |
| | Conductor (metal) | 2 $\mu\text{m}$ | 1 $\mu\text{m}$ |
| **Multi-layer** |  | No | Yes |
| **Other materials** |  | No | Yes |

To further promote access, a set of “off-the-shelf” pMEAs were designed, fabricated, packaged, tested, and stocked. These are available for users who want to try the technology and do not need custom solutions. Example designs include a 64-channel penetrating pMEA for rats and small mammals, described previously and dubbed the PIE Foundry ‘standard’ array (Fig. 2) [58], and a 32-channel penetrating tetrode array.

**Figure 2.**
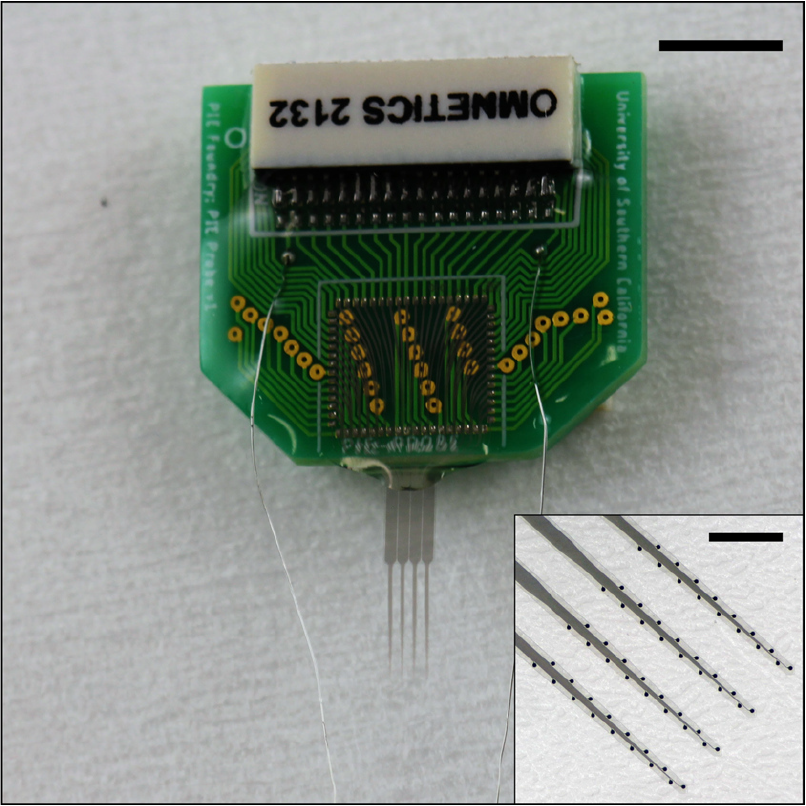
Photograph of the PIE Foundry ‘standard’ penetrating microelectrode array. Scalebar is 5 mm. The polymer-die is ultrasonically welded to a custom circuit board with two back-and-front Omnetics brand analog connectors, and a pair of soldered stainless steel ground leads. Inset shows magnified view of the platinum coated electrode sites, 30 µm in diameter. Inset scalebar is 1 mm.

The PIE Foundry uses an online portal and direct emails for receiving user-submitted digital design files and proposals [59]. Microfabrication services typically take 1-2 months depending on the current workload, while custom projects can take 3-5 months following scientific steering group approval, depending on complexity of the project. Users submit a standard user agreement, outlining intellectual property guarantees and confidentiality of submitted designs, as the only requirement for access. Designs for pMEAs that interface to the brain, peripheral nerves, spinal cord, and retina have been produced for users around the world.

### 2.2. Fabrication methods

Microfabrication of pMEAs uses a combination of photolithography, physical vapor deposition, and plasma etching (Fig. 3). This process produces bare arrays that then need to be packaged to establish connections to electrophysiological recording equipment and/or encapsulated to provide protection against water or saline intrusion into electrical contacts between the arrays and external connections.

**Figure 3.**
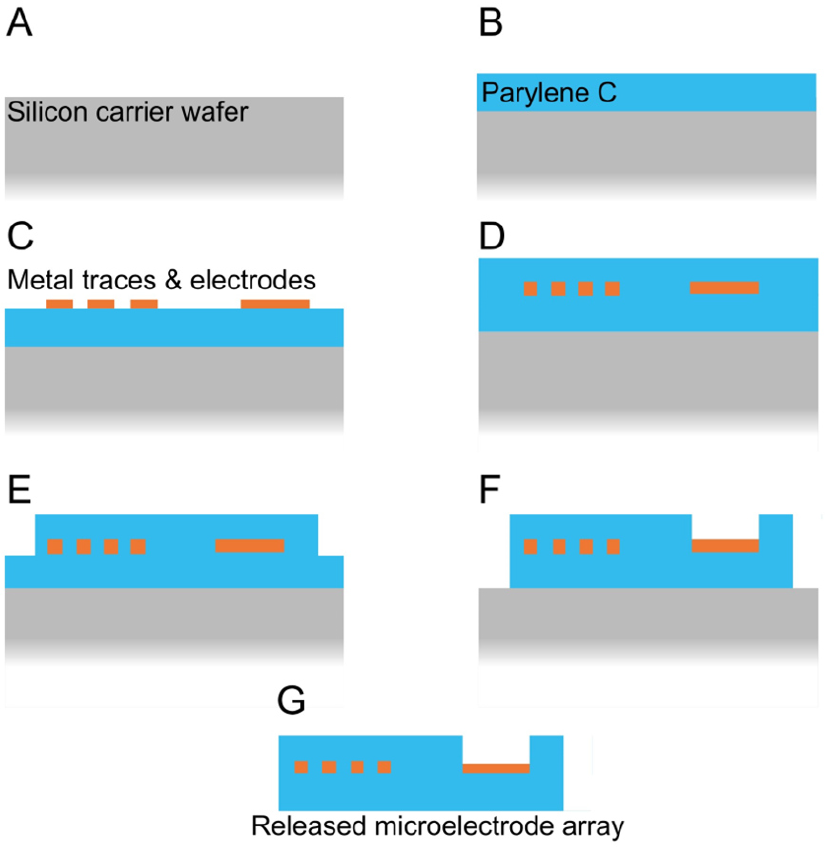
Depiction of the cross-section of a polymer microelectrode array during typical fabrication: **A** Silicon carrier wafer; **B** Parylene C base layer coating; **C** Metal traces and electrodes deposited with evaporation and defined with lift-off lithography; **D** Parylene C insulation layer coating; **E** Outline of array partially etched with oxygen plasma; **F** Array is cut-out and electrodes exposed with second oxygen plasma etch; **G** Array is released from carrier wafer.

All processing is performed on 100 mm diameter silicon carrier wafers. Larger or smaller wafers can be used, and the size is dictated by the available process equipment. In contrast to semiconductor integrated circuit foundries that utilize large diameter wafers (200 or 300 mm) to increase throughput and decrease cost, university-based facilities that conduct research are still largely limited to 76, 100, or 150 mm substrate diameters. Such wafer sizes are cost-effective and sufficient in usable area for low to medium volume manufacturing of pMEAs.

Untreated silicon wafers possess a thin layer of native oxide which facilitates later release of the pMEAs due to the weak adhesion of Parylene to SiO_2_. The silicon carrier wafer provides a convenient starting point for processing as wafers are widely accessible with high-quality, flat, and smooth surfaces. They are also relatively inexpensive ($15-20/wafer in small volumes).

Full details of the fabrication have been made available in an open-source repository [60]. All pMEA fabrication is performed in a class ISO 6 cleanroom. First, carrier wafers are coated in approximately 10 µm of Parylene C to form the base layer of the pMEA (PDS 2010, Specialty Coating Systems, Indianapolis, IN). Since Parylene coating is conformal, this process coats all exposed wafer surfaces, including the backside and edges. This prevents the processed polymer film from separating from the carrier wafer during the subsequent steps. Wafers are then baked in an oven under vacuum or a nitrogen atmosphere at 150 °C for 4 hours, which increases the crystallinity of Parylene, decreasing its thermal expansion in preparation for the subsequent metal deposition step.

Next, a single thin-film metal layer, containing all traces, contact pads, electrodes, labeling, and markings, is fabricated using a combination of image-reversal photolithography and solvent facilitated metal lift-off. The lithography step begins by dry-baking Parylene C coated wafers for 30 minutes in a vacuum oven at 60 °C and 15” Hg, with approximately 50 sccm of nitrogen flow. This step helps improve adhesion between the photoresist and Parylene C surface. Wafers are spin coated with 3 mL of de-gassed AZ5214 photoresist (Integrated Micro Materials, Argyle, TX) in a two-step process (500 RPM 5 seconds; 3200 RPM 40 seconds; Laurell Technologies model WS-400, Landsdale, PA). The final thickness is 1.15 ± 0.1 µm. Photoresist coated wafers are soft-baked at 110 °C for 60 seconds to remove residual solvent and are then exposed to ultraviolet (UV) radiation through a chrome photomask in hard-contact mode at 42 mJ/cm^2^ on a contact aligner. Wafers are baked at 110 °C for 63 seconds to initiate the image-reversal mode of AZ5214, then allowed to rest for at least 3 minutes. This second bake step is critical and very sensitive to small changes in temperature; hotplate surface temperature and its uniformity across the surface is confirmed with an infrared thermometer (Fluke, Everett, Washington). Wafers are then UV-exposed over the entire surface, without a photomask, at 280 mJ/cm^2^, and immediately placed in a room temperature deionized (DI) water bath for 2 minutes. The pattern is developed by submerging wafers in a developer bath (1:4 mixture of AZ340k in DI water) for 18 seconds with mild agitation, followed by serial rinses in DI water. In preparation for metal deposition, wafers are exposed to a mild O_2_ plasma (100W, 100 mT, 300 seconds, YES CV200 RFS, Fremont, CA) to roughen the Parylene C surface and improve adhesion.

The metal layer is deposited by e-beam evaporation using a CHA Mark-40 electron-beam evaporator (CHA Industries, Fremont, CA). Typical processing entails either a single 200 nm Pt layer or a Ti/Pt/Au/Pt stack of 20/25/155/25 nm. The patterned metal-layer is defined using solvent lift-off to remove the un-patterned photoresist. Lift-off is performed in a bath of N-methyl-2-pyrrolidone (NMP) at 60 °C. Alternatively, wafers can be soaked overnight in acetone at room temperature. Pt coated wafers are treated with mild agitation while Ti/Pt/Au/Pt coated wafers require an ultrasonic bath. The process takes approximately 10-20 minutes. Wafers are then cleaned in serial rinses of room-temperature NMP, isopropanol, and DI water.

A second Parylene C layer, also approximately 10 µm thick, is deposited to insulate the metal layer. Wafers are first dry-baked for 30 minutes at 60 °C and 15” Hg of vacuum, under 50 sccm of N_2_ flow, and are again treated with O_2_ plasma to roughen the Parylene C base layer and improve adhesion. Total thickness of the multi-layer stack is approximately 20 µm. After depositing the second Parylene C layer, the entire wafer is typically annealed a second time under vacuum or nitrogen atmosphere for 4 hours at 150 °C. This step helps prevent mismatch in the mechanical properties between the two layers of Parylene C.

In order to expose the electrodes and contact pads and to cut-out the shape of the pMEA, wafers are etched with O_2_ plasma through two separate photoresist masks. The first mask contains the outline of the MEA, while the second mask contains the outline, as well as the shape of the exposed contact pads and electrodes. Wafers are spin coated with 3 mL of de-gassed P4620 photoresist (Integrated Micro Materials, Argyle, TX) in a two-step process (500 RPM 5 seconds; 1000 RPM 45 seconds). The final thickness is 15±1 µm. Photoresist coated wafers are soft-baked at 100 °C for 20 minutes to remove residual solvent. The edge-bead is manually removed using a swab soaked in edge-bead removal solvent. Wafers are then aligned to a chrome photomask using the visible metal layer for alignment and are then UV-exposed in vacuum contact at 480 mJ/cm^2^. Wafers are immediately quenched in room temperature DI water bath for 3 minutes to prevent bubbles forming in the Parylene C from the exothermic photo-initiated reaction [45]. The pattern is developed by submerging wafers in a developer bath (1:4 mix of AZ340K to DI water) for approximately 2-3 minutes with mild agitation, followed by serial rinses in DI water.

10 µm of Parylene C is etched using the first photoresist mask in either an O_2_ RIE process or a switched-chemistry Bosch-like process [61]. O_2_ RIE is performed using an Oxford RIE80 (Oxford Instruments, Abingdon, United Kingdom) at 150 mT, 150 W, 50 sccm of O_2_, yielding an approximate etch rate of 0.22 µm/minute and an approximate selectivity of 1:1. The switched chemistry process also uses O_2_ but in a deep reactive ion etcher (DRIE) typically used for high aspect ratio etching of silicon (Oxford RIE-STS, Oxford Instruments, Abingdon, United Kingdom). In a two-step switched-chemistry process, alternating, short-duration O_2_ etch (700 W ICP power, 20 W RF power, 60 sccm O_2_, 40 sccm Ar, 10 seconds) and C_4_F_8_ passivation steps (700 W ICP power, 10 W RF power, 35 sccm C_4_F_8_, 40 sccm Ar, 3 seconds) are repeated until the process endpoint, yielding an approximate etch rate of 0.085 µm/cycle (0.40 µm/minute). Typically etching is performed in batches of 25-30 cycles to prevent photoresist hardening from the combination of heat, UV, and plasma exposure. Following the first etch step, the first etch mask is stripped in acetone, cleaned in isopropanol and DI water, and then baked dry. The second etch mask is applied in the same manner, and the etching process is repeated. The second etch mask is then stripped, and the devices are released by submerging the wafer in deionized water and peeling devices off the wafer using tweezers.

Devices are annealed for 48 hours under vacuum at 200 °C to improve adhesion between the Parylene C layers. PMEAs are typically sandwiched between ceramic plates (0.025” thick) to reduce curvature [62]. After annealing devices are treated with a brief exposure to O_2_ plasma (100 W, 100 mT, 300 seconds; YES CV200 RFS).

### 2.3. Multi-project wafer processing

Submitted designs which adhere to a standard set of guidelines can be processed as part of MPW services. All MPW fabrication incorporates a 10 µm base layer of Parylene C, a 10 µm insulation layer of Parylene C, and a single 200 nm metal layer. Guidelines for MPW processing dictate a minimum metal feature resolution of 2 µm (4 µm pitch), a minimum resolution of etched features of 10 µm, and an alignment tolerance between lithography layers of ± 5 µm. MPW processing is performed on 100 mm Si carrier wafers, with 5 mm edge borders, allowing approximately 6350 mm^2^ of available space. Between 3 and 6 user designs are typically allocated depending on demand and device size. MPW designs are submitted from multiple users as digital files (.gbr, .dwg, .dxf, or .gds) through a webform, and combined into a single set of photomasks. Because Parylene C devices are released through etching instead of dicing, devices regardless of shape can be tessellated without relying on a regular grid layout, increasing density, and minimizing dead-space.

PMEAs are typically panelized with tab-routing (Fig. 4), a practice borrowed from the PCB manufacturing industry. The cut-out etch mask is modified such that devices from a single user are released in a continuous panel, with the individual pMEAs connected via 2-3 slender Parylene C tabs, typically 1-2 mm in length and width. Panelizing pMEAs in this manner allow arrays to be post-processed, handled, stored, and shipped as a single unit. Users cut out individual devices at the tabs, which then provide a place for tweezers to grip the pMEA without damaging the electrodes or contacts. Panels are stored and shipped sandwiched between layers of either conductive polyethylene plastic or high-density polyester fabric (Texwipe TX1004, TestEquity, Moorpark, CA), which are then sandwiched between squares of 1/8” corrugated plastic. The corrugated plastic serves to keep the pMEAs flat during storage and transit, while the inner layer prevents the pMEAs from moving due to static buildup.

**Figure 4:**
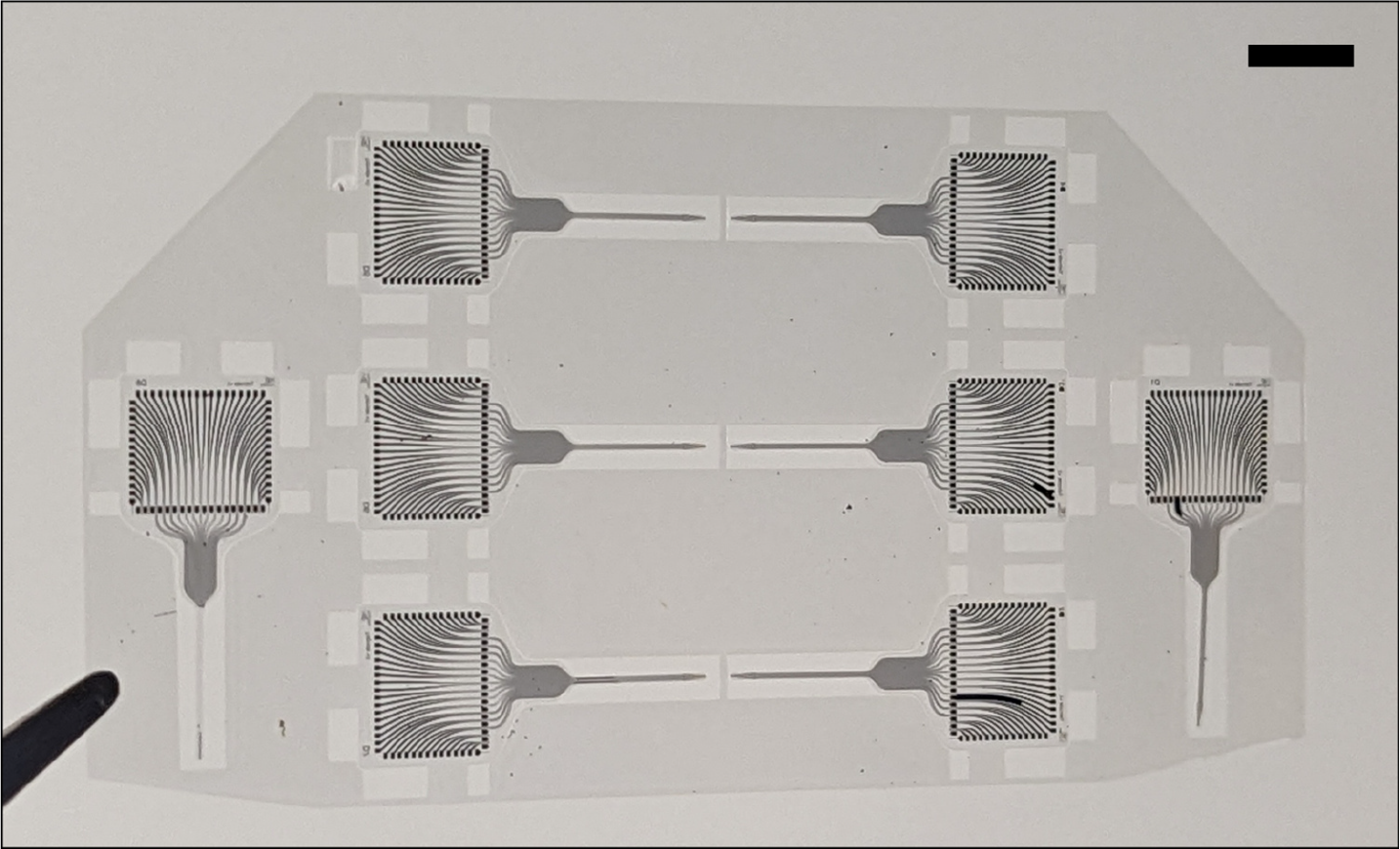
Photograph of panelized polymer microelectrode arrays with tab-routing. Scalebar is 5 mm.

MPW runs are completed every 4-6 weeks and are scheduled every 1-2 months depending on demand. A typical MPW run includes 6-12 100 mm diameter wafers, and the number of possible pMEAs is dependent on the size, as larger arrays demand more space and therefore fewer can be fabricated. The number of completed pMEAs per user ranges from as few as 5 to as many as 200.

### 2.4. Packaging and PCB connection

The majority of devices manufactured by the Foundry are attached to PCBs for connection to external electronics and headstages, and for connection to macro reference and ground electrodes. Connection to PCBs is typically achieved with a direct ultrasonic welding method referred to as polymer-ultrasonic on bump (PUB) bond [63]. PUB bonding is a simple and effective method to attach pMEAs to PCBs without the need for solder, conductive films, or zero-insertion force (ZIF) connectors. This approach helps minimize both the size of the PCB and the contact pads on the pMEA, scales easily to pMEAs with high channel counts (>100), avoids the use of non-biocompatible materials, can be performed very quickly (approximately 2 seconds per connection), and uses commonly available ball-bonder type wire-bonding tool.

PUB bonding contact pads are typically 200 × 350 µm^2^; pMEA pads have been fabricated from Au, Pt or Au/Pt stacks, while the corresponding pad on the PCB consists of 1 oz Cu with a finish of soft, bondable gold 20-30 microinches thick. Pads are laid out as a rectangular grid or along a rectangular perimeter depending on space available for trace routing.

PCBs are first ‘bumped’ using a ball-bonder type wire-bonding tool (Hybond Inc. model 626, Escondido, CA). A gold ball with an approximate diameter of 100 µm is ultrasonically bonded to the center of each pad. Each ball is then flattened into a disk using a wire-bonding tool with a tape-automated bonding (TAB) attachment (7045-Ti, Smart Precision Tools). This step flattens and levels the gold bump, while imprinting a waffle pattern onto the surface. The pMEA is then aligned to the PCB contacts using a stereoscope and temporarily held in place with any common adhesive tape. Using the same wire-bonder tool with the TAB attachment, force and ultrasonic energy is applied to the back of the pMEA, welding the metal of the pMEA contact pad to the gold bump. Each pad is bonded, the process typically takes just 2 seconds per pad. Afterwards the bonded connection is underfilled with epoxy (301-3M, EpoTek) and baked on in an oven for 1 hour at 55 °C, after which the tape is removed. pMEAs are typically then overcoated in a thicker epoxy (302-3M, EpoTek) to provide encapsulation and protection.

When required, PUB bonding can be replaced with alternative packaging schemes, including the use of ZIF connectors. ZIF connectors offer reversible connection between flat flexible cables and PCBs using a mechanical clip. ZIF compatible contact pads are incorporated into the pMEA design, oversized by 2.5% to account for polymer shrinkage during annealing steps [42]; pMEAs are then mounted onto backer films, typically either PEEK or polyimide films of a thickness specified by the manufacturer of the ZIF connector (typically 250 µm), using a non-viscous epoxy (e.g. EpoTek Technology 301 epoxy, Billerica, MA). The film provides the necessary thickness and mechanical support to mate with the ZIF connector.

In all cases, PCBs and components are encapsulated in 10 µm of Parylene C prior to packaging the pMEA, as a means to prevent moisture intrusion leading to crosstalk between channels. Thin strips of adhesive tape are used to mask important areas such as connector faces and bond pads, then removed using a razor blade after Parylene C coating. PCBs are typically coated in a thick layer of biocompatible epoxy as a final step.

### 2.5. Characterization

Representative electrodes from each fabrication run are characterized with electrochemical impedance spectroscopy (EIS) and cyclic voltammetry (CV) using a Gamry Reference 600 potentiostat (Gamry Instruments, inc., Warminster, PA). CV is performed in 0.05 M H_2_SO_4_ in the potential range of −0.2 - 1.2 V versus an Ag/AgCl reference and large Pt counter electrode. The solution is purged with N_2_ prior to and during the CV. The typical scan rate is 250 mV/s for 30 cycles. EIS is performed in 1× phosphate-buffered saline (PBS) following CV cycling, also using Ag/AgCl and Pt as the reference and counter, respectively. Impedance is measured in the range of 1 Hz–1 MHz. However, unless otherwise requested, pMEAs shipped to users are only characterized using single-point electrochemical impedance measurements as a means to assess end-to-end continuity and confirm electrode qualities do not vary significantly from expectation. Impedance is measured at either 1 or 10 kHz in 1× PBS against a Pt counter. Electrodes are tested for open-circuits, shorts, or excessively high impedance. In addition, pMEAs are examined under confocal and stereomicroscopy for defects, including delamination of contact pads or electrodes, cracks in the polymer base or insulation, and cracks in the metal microscale traces.

### 2.6. In Vivo validation

Implantation strategies for PIE Foundry pMEAs vary with target anatomy, and pMEA size and design, and are typically devised by the user based on the needs of their experiment. Flexible pMEAs targeting brain regions deeper than 1-2 mm typically require either a shuttle or temporary stiffening technique [53,58]. A simple protocol using dip-coated bio-dissolvable polyethylene glycol (PEG) was developed to provide users with a generalizable approach. The technique was designed to require minimal equipment or materials, and to minimize the acute displacement of brain tissue during the surgery.

During preparation PEG with 8 kDa molecular weight is melted in a microwave or on a hotplate. The PEG is briefly heated to just above 100 °C then allowed to cool to between 60 and 80 °C. The molten PEG is kept in a small beaker or petri dish on a hotplate at a constant temperature. The pMEA is soaked in 70% ethanol to aid in sterilization and fully dried, then attached to a micro-manipulator which can smoothly raise or lower the assembly. The array is lowered into the PEG until the bottom edge of the PCB is in contact with the liquid, then the device is slowly retracted at a speed of approximately 20 µm/sec. The thickness of the PEG coating can be adjusted by maintaining the PEG solution at different temperatures or adjusting the rate of withdrawal. Lower temperature and slower retrieval speed can generate thick, brace-like PEG coating, while a thinner coating can be generated by using PEG at higher temperatures. Due to surface tension effects, bridging PEG films can form between Parylene C shanks on arrays with multiple shanks. If this occurs the array is briefly submerged in pre-sterilized distilled water, only to the depth of the shanks, for 30 seconds to 2 minutes, dissolving the PEG film. After fully drying, the Parylene C shanks are lowered into the PEG solution to re-coat.

To validate the *in vivo* performance of pMEAs built using our approach, examples of the PIE Foundry 64-channel ‘standard’ array were implanted into the hippocampus of adult Sprague-Dawley (SD) rats. All procedures followed the Guide for the Care and Use of Laboratory Animals and approved by the University of Southern California Institutional Animal Care and Use Committee. The implantation procedures are detailed in our previous conference proceedings [58]. In brief, a 2x4 mm^2^ bone window above the right hippocampus is drilled open. Both the dura and pia layer are carefully removed to expose the brain surface. The PEG-coated pMEA is slowly inserted with a micro-manipulator at a speed of 100 µm/second to the desired depth. Ground wires are inserted into a small hole drilled above the cerebellum. For chronic implantation, the pMEA is fixed onto the skull by attaching it to five anchor screws pre-secured on the skull. After 3 to 7 days of recovery, neural activities are recorded with a 64-channel data acquisition system (Plexon Inc, Dallas, Texas) while the animal is free exploring in a round open field. Broadband signals are high-pass filtered for better identification of spike activities. Food pellets are randomly scattered on the floor of the open field to encourage the continued exploration of the animal. After all recordings are taken, the animal is euthanized, and the brain tissue is preserved. The location of the implanted pMEA is verified through histology.

## 3. Results

### 3.1. Shared resource model

The PIE Foundry launched in November 2019 and has partnered with more than two dozen labs worldwide to deliver more than 500 pMEAs of varied design, application, target anatomy and animal model. Microfabrication of a pMEA from a submitted design can be accomplished in as quickly as one month, while custom projects, including pMEA design and fabrication, PCB design and purchase, packaging, and testing, are typically accomplished in 3-5 months. These turnaround times are short, even compared to commercial entities offering custom fabrication runs (based on our own experiences and conversations with our users, custom silicon MEA orders may take upwards of 6-8 months for delivery) and amounts to a significant time savings compared to the duration required to train a graduate student and the commitment to develop such technology in an academic research group. We have been able to assist users from four different continents, covering researchers at a broad range of career-stages and institutions, while maintaining a zero-cost fee structure for non-profit users.

### 3.2. Fabrication & packaging

The fabrication methods described above have been demonstrated as flexible and robust; using this single set of techniques we have built peripheral nerve interfaces, surface arrays, and penetrating arrays, targeting animal models including songbird, mouse, rat, cat, and sheep, with channel counts spanning 2 to 64. Examples of manufactured electrode layouts include stereotrodes, tetrodes, linear arrays, grids, and anatomically mapped custom arrays, for electrodes ranging in size from < 200 µm^2^ to 1,000,000 µm^2^. Users have successfully deployed our pMEAs for spike sorting and tracking of individual units, as well as measuring local field potentials, and stimulating peripheral nerves and spinal nerves. pMEAs have been used for acute experiments, and free-moving chronic experiments lasting several months. Moreover, we were able to build arrays of such varied design and size simultaneously, on a single wafer, as part of our MPW fabrication runs. When engaging a custom fabrication project, we have been able to modify our fabrication techniques to incorporate features including through-layer vias, for multi-layer pMEAs, thermoformed 3D shapes, and exotic materials including glassy-carbon electrodes, or nanostructured coatings.

These lithographic methods allow reliable micromachining of critical features, including metal traces, with size and spacing as small as 2 µm. Successful device yield has reached >80% as of 2023, as determined by a threshold of >95% channel functionality, and <5% variation in electrode impedance and critical dimensions from design.

The PUB bonding method has proven the most effective packaging method, more scalable and more robust than the use of ZIF connectors, and faster and more reliable than manual alternatives such as conductive epoxy. A 64 channel pMEA can be PUB bonded to a PCB in just a few minutes, requiring a space one third the area of the smallest available corresponding ZIF connector. The technique easily scales to hundreds of channels, far more than the largest available ZIF. Across more than 100 tests, we have yet to observe a PUB bond fail after packaging.

Figure 5 presents representative datasets showing electrode CV response and electrochemical impedance. Electrode impedances scale inversely with electrode size. While the PIE Foundry does not offer electrochemical coatings, several users have used PIE Foundry devices with commercial or custom systems for coating porous gold, PEDOT, and other surface finishes, and have achieved significant decreases in impedance.

**Figure 5:**
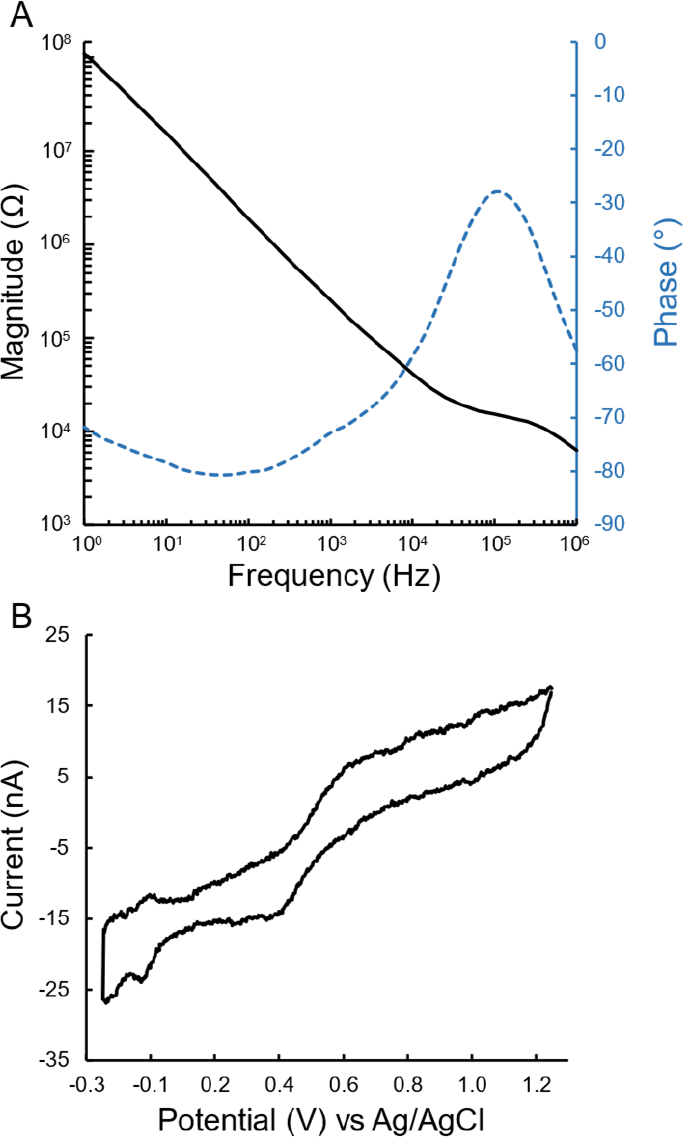
Representative examples of **A** electrochemical impedance spectroscopic and **B** cyclic voltammetry data for a 30 µm diameter Pt coated microelectrode from a PIE Foundry hippocampal polymer microelectrode array.

### 3.3. In Vivo validation

We performed eleven implantations of the PIE Foundry ‘standard’ array. All pMEAs were dip-coated following the protocol described above. In two of the implantation surgeries the pMEA shanks deviated from the target, with shanks deflecting off tissue at a depth of 2 mm. The other nine implantations were all successful; electrodes were implanted into rat hippocampi smoothly and without complication. Histological micro-images of brain slices collected after ending experiments (Fig. 6) were used to verify placement of the electrodes, and to confirm that the shanks remained straight during implantation.

**Figure 6.**
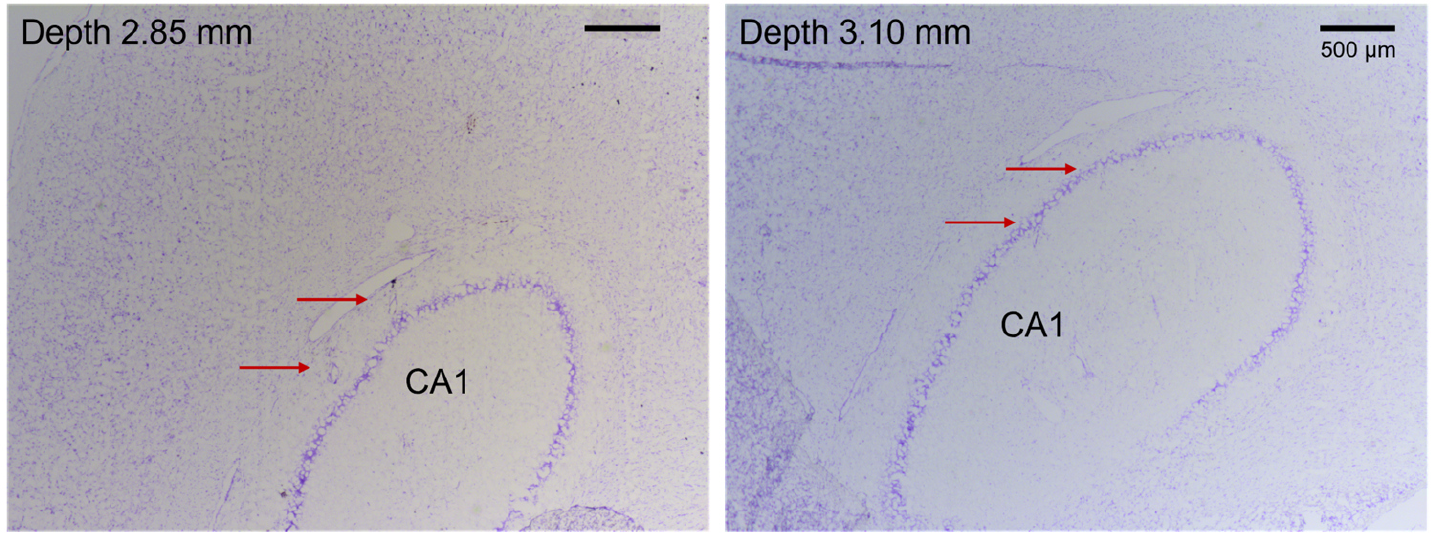
Microimages of 50 µm thick, Nissl-stained coronal brain slices collected from one animal implanted with a 64-channel Parylene C based microelectrode arrays, one month after implantation. Slices are collected at a depth of 2.85 mm (left) and 3.10 mm (right). Red arrows indicate tracks left by the implanted array.

Neural activities were monitored and recorded during and after implantation. After the pMEA was inserted into the desired depth, up to 44 units were recorded from an individual anesthetized animal. Complex spikes, which are the signature firing pattern of hippocampal pyramidal neurons, were recorded along electrodes up and down the Parylene C shanks. These recordings demonstrated that the standard pMEA is capable of recordings which span multiple hippocampal cell body layers and obtaining neural activities from multiple sub-regions. After recovery neural signals were recorded from three chronically implanted animals. Up to 62 units with signal-to-noise ratio (SNR) > 3 were recorded four days post-implantation (Fig.7), and 36 units were recorded one-month post-implantation from one animal. Spike activities with stable waveform and amplitude were recorded from two channels over a duration of 34 days from another animal (Fig. 8). These preliminary chronic recordings demonstrate that the standard pMEA can obtain high-quality unitary activities from free-behaving animals. Although we have not conducted strict analysis to track individual neurons over time, stable recording of spikes with similar features indicates that the surrounding tissue remains healthy, and the electrical properties of the electrodes are stable unde*r in vivo* conditions. No damage to the packaging was noticed for the chronically implanted period which demonstrated that the package is robust and can sustain the daily behaviors of the animal such as scratching and bumping into the cages.

**Figure 7.**
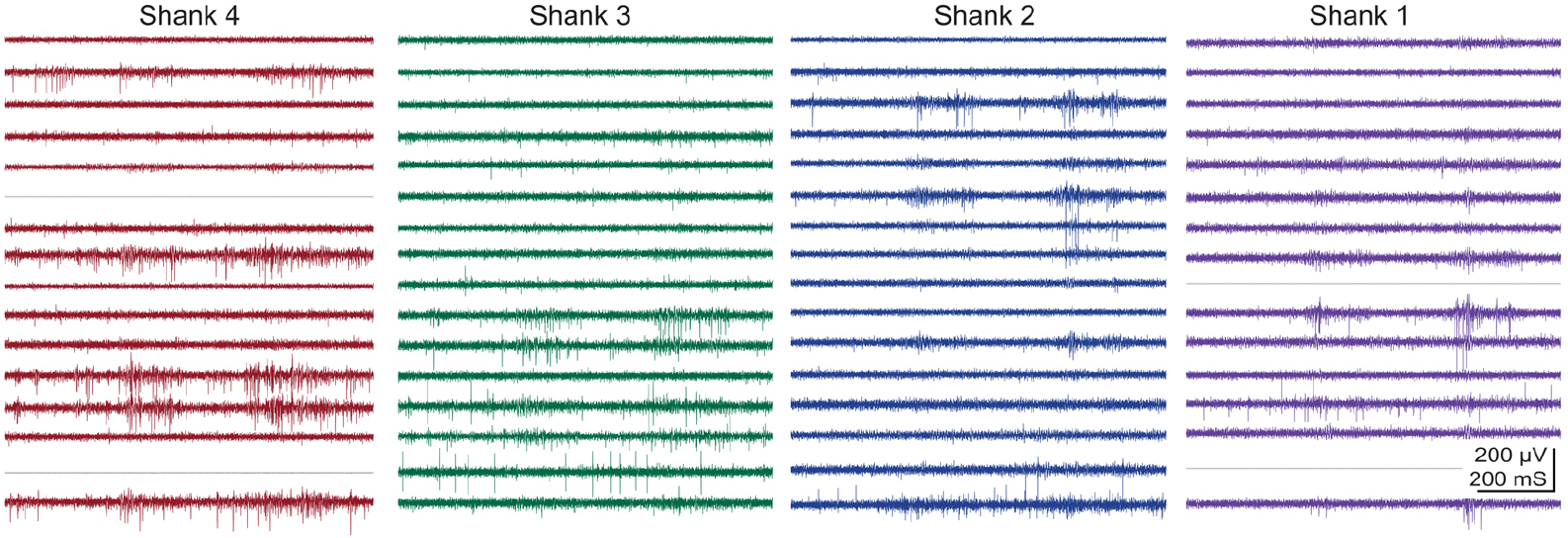
Representative data recorded from rat hippocampus using a 64-channel Parylene C based microelectrode array. Signals were recorded four days post implantation and high-passed filtered. All 64 channels are presented.

**Figure 8.**
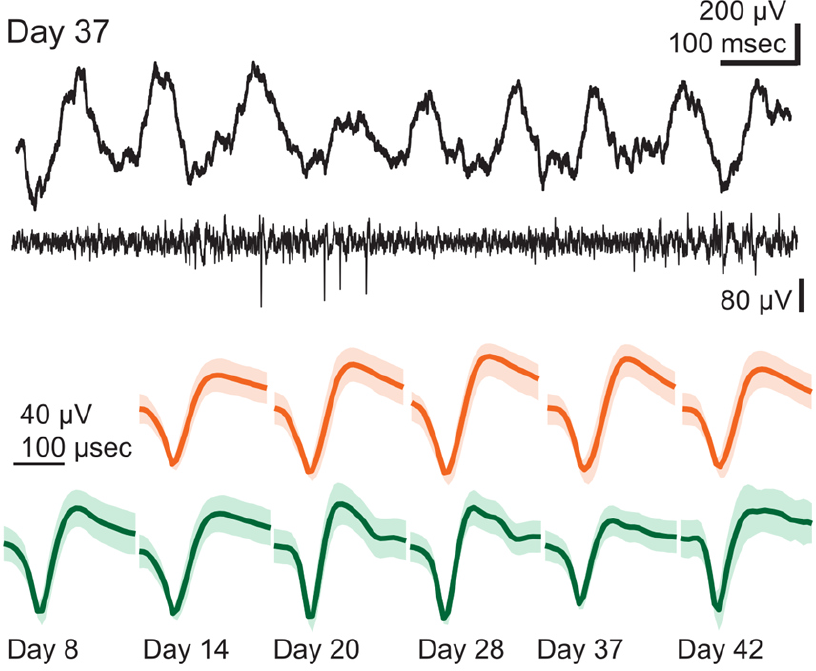
Representative data recorded from rat hippocampus using a 64-channel Parylene C based microelectrode array. (Top) Local field potentials and spikes recorded 37 days post-implantation. (Bottom) Individual neuronal units with stable waveforms and amplitudes recorded on two different channels (orange and green) over 34 days.

## 4. Discussion

The *in vivo* results presented here demonstrate that our pMEAs are capable of resolving unitary activities from neurons in free-moving animal models with high signal-to-noise ratios, and the preliminary chronic recording data indicate our pMEAs can obtain stable, long-term recording. These results are specific to the PIE Foundry ‘standard’ array, depicted in Figure 2, and the experiments described above, but reports from users indicate similar success using a wide range of designs in a wide range of animal models. Likewise, users have reported successful deployment of PIE Foundry fabricated pMEAs for use in peripheral nerve stimulation and surface recordings. Detailed results from our users have been presented at conferences [58,64,65] and are being compiled into peer-reviewed articles under the independent authorship of each user institution [66,67]. The nature of the shared-resource model is that the data collected by PIE Foundry users remain the intellectual property of the user, who retains publication rights. Collecting feedback on electrode performance is typically conducted with follow-up interviews with users and used to address shortcomings or create revised versions of previously manufactured arrays. Many users have become repeat customers, developing several generations of pMEA designs.

Nearly four years after the launch of the PIE Foundry, demand for our pMEA manufacturing services continues to increase. Users cite a lack of alternatives. Despite the growing role of flexible electrodes in the development of brain-computer-interfaces there remains few avenues outside of academic cleanrooms to supply such devices, especially for researchers who need custom solutions. At the same time the technology has matured to a point where there are fewer opportunities for graduate students to generate novel, publishable data on the microfabrication of pMEAs; this conflict between the interests of engineering students and neuroscience researchers, and the dearth of commercial options, has highlighted the on-going need for shared-resource centers like the PIE Foundry.

Four years of operation have also revealed the major obstacles facing all engineers attempting to advance pMEA technology. Many of these are technical challenges in the microfabrication and packaging of electrodes, which we believe the methods described here largely resolve. Another is the challenge of scaling up channel counts while managing analog outputs; many researchers are interested in increasing pMEA channel counts into the hundreds or shrinking the size of the electrode interface boards while maintaining the same channel counts for recording from smaller animal models. In both cases, the need for bulky analog connectors is a major limiting factor. Integration of pMEAs with multiplexing circuitry has been accomplished [6,21], but this approach relies upon Intan Technologies products which only reach 64 channels and impose a high component cost. Other major engineering challenges include the appearance of channel crosstalk in some chronically implanted pMEAs (> 1 month), which we have been able to resolve with appropriate thermal annealing steps and water-proof encapsulation of the electrode interface board; Both processes were described above.

The most significant outstanding issue concerns implantation and user training. There remains no agreed upon best-practice for implanting a flexible pMEA into neural tissue. The above-described PEG dip-coat method was chosen because it is simple to perform and requires no special equipment, and therefore can be easily prepared by users. But the method requires the removal of the pia and dura mater in most surgeries, which can cause damage to superficial cortical areas. If the coating is prepared improperly, the shanks of penetrating array may follow a curved path, making accurate placement difficult. These drawbacks can be addressed by using an implantation shuttle method, temporarily attaching a microneedle to the polymer [57,68–70]. But shuttle approaches require tedious and difficult manual manipulation of the probe, they scale poorly to large numbers of arrays, they are difficult to use with multi-shank arrays, and many involve the use of a bioresorbable adhesive which can be incompatible with sterilization methods. Developing a simple and generalizable implantation method would be a pivotal step in expanding the accessibility of pMEAs. Another barrier is the limited or lack of experience with implantation of flexible probes. Users, often graduate students, require training in the handling and use of pMEAs, in addition to training on surgical technique and implantation methods. In response, the PIE Foundry launched a user training service, which includes an annual series of free, online workshops, and more recently an offering of free in-person training. Eight workshops have been held since 2020, with more than 130 trainees in attendance from 59 different institutions and 12 countries. In-person training began in 2023, with two sessions held so far. These sessions include presentations on pMEA design and fabrication, implantation, including craniotomy and dip-coat preparation, and application, including data collection from a free-moving animal model. Finding scalable ways to train researchers and surgeons on the use of pMEAs will be critical for expanding access to this technology.

Ultimately for pMEAs to undergo widespread adoption, some form of commercialization is likely required, whether this entails the sale of a limited set of commercial pMEAs or design and fabrication services from commercial facilities, mimicking the path followed by silicon MEAs. A steady, recurring demand for polymer electrodes necessitates a trained user community. Until then, publicly funded service centers like the PIE Foundry will fill a critical role in the development and dissemination of this technology.

## 5. Conclusion

A shared-resource center was established for manufacturing polymer-based microelectrode arrays for use as neural interfaces in research. A comprehensive set of scalable and generalizable microfabrication processes were developed and validated for batch-scale production of pMEAs of arbitrary user-submitted custom designs. Processing capabilities include pattern resolution down to 2-micron feature size, successful yields greater than 80%, and batch production of 10s to 100s of pMEAs on a monthly basis. Additional protocols and techniques were developed for packaging, characterizing, implanting, and using pMEAs, and disseminated through online and in-person training sessions. Since 2020, more than 500 fully functional pMEAs have been delivered to more than two dozen separate labs, with designs including peripheral nerve interfaces, surface arrays, and penetrating arrays, targeting animal models including songbird, mouse, rat, cat, and sheep, with channel counts spanning 2 to 64, for applications including both neural recording and stimulation.

## Acknowledgements

This work was funded through the BRAIN Initiative and the National Institute of Health under award U24NS113647. The authors would like the thank the staff at the John O’Brien Nanofabrication Facility at USC for assistance.

